# HOMED enables hierarchical and multimodal optimization of DNA methylation deconvolution across tissues

**DOI:** 10.64898/2026.06.08.730896

**Authors:** Yuansen Liu, Yundi Chen, Yuheng Du, Lana X Garmire

## Abstract

Cellular heterogeneity is a major confounder in bulk DNA methylation data used in epigenome-wide association studies. Existing reference-based DNAm deconvolution methods often overlook hierarchical relationships among related cell types and may generalize poorly across datasets due to limited variability in reference profiles. We developed **HOMED (Hierarchically Optimized Methylation Deconvolution)**, a framework that integrates cell-type hierarchy, single-cell RNA sequencing-guided deconvolution, and paired bulk RNA-seq/DNAm data for CpG signature optimization. Across multiple simulated and real heterogeneous tissue datasets, HOMED achieved a 68% higher mean Pearson correlation coefficient (PCC) and 60% lower mean root mean squared error (RMSE), compared to other scRNA-seq-guided or single-cell DNA methylation-based DNAm deconvolution methods. Moreover, application of HOMED to a smoking-exposed placenta DNA methylation cohort revealed significant shifts in cell-type compositions in endothelial, nRBC, stromal, and villous trophoblast proportions. Adjusting for these compositional differences reduced inflated smoking-associated CpGs by nearly three folds, and doubled the number of enriched pathways with known associations to smoking. In summary, HOMED is an accurate and practical method to infer cell-type proportions from bulk DNA methylation data.

**Graphical Abstract:** 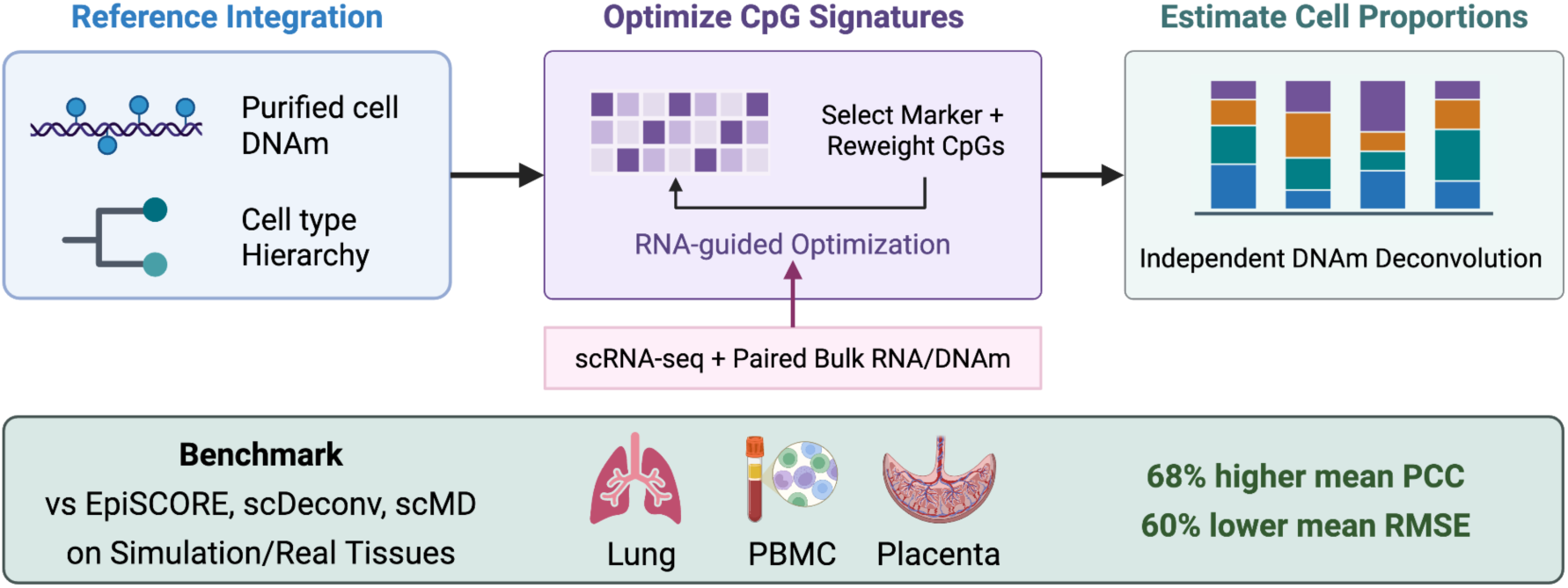

## Introduction

Bulk tissue samples are inherently heterogeneous, and variation in cellular composition represents a major source of confounding in epigenomic studies. In DNA methylation (DNAm) analyses, differences in cell-type abundance can obscure biologically meaningful signals and produce spurious associations if cellular heterogeneity is not appropriately accounted for (1). Therefore, accurate deconvolution to estimate cell-type proportions from bulk DNAm data is a critical preprocessing step in translational epigenetics research.

Reference-based approaches remain the dominant strategy for DNAm deconvolution, modeling bulk methylation profiles as mixtures of cell-type-specific signatures. Traditionally, these signatures have been constructed directly from purified cell populations, such as in Houseman’s method (2). More recently, single-cell datasets have provided complementary information for constructing or refining cell-type-specific DNAm references. For example, EpiSCORE (3) leverages single-cell RNA sequencing (scRNA-seq) data to derive cell-type-specific gene expression profiles and uses the relationship between gene expression and DNA methylation (DNAm) to infer cell-type-specific methylation signatures. Similarly, scDeconv(4) exploits scRNA-seq-derived cell-type information to construct reference profiles for estimating cellular compositions from bulk DNAm data. On the other hand, scMD(5) directly utilizes single-cell DNA methylation (scDNAm) data to characterize cell-type-specific methylation patterns and construct reference profiles for DNAm deconvolution, thereby avoiding reliance on cross-omic relationships between gene expression and methylation. These approaches highlight the potential of leveraging single-cell data to improve reference construction and cell-type resolution in DNAm deconvolution, while illustrating distinct strategies based on cross-omic inference versus direct methylation profiling.

Despite these advances, many limitations remain. First, most methods treat cell types as independent relationships and do not explicitly model hierarchical relationships among related lineages and subtypes. Since closely related cell types often exhibit highly similar DNAm profiles, ignoring hierarchical structure may reduce deconvolution accuracy at a fine resolution (6). Recent advances in single-cell and single-nucleus RNA sequencing have enabled increasingly refined cell-type reference atlases, creating new opportunities for hierarchy-guided DNAm deconvolution. Second, existing reference optimization algorithms generally optimize a common CpG library across all cell types. For example, IDOL iteratively identifies CpGs that maximize overall deconvolution accuracy, with performance evaluated across cell types (7). However, a globally optimized library may not be equally informative for every cell population, particularly when cell types are closely related. Cell-type-specific optimization may therefore be preferable, allowing CpG libraries to be optimized according to their ability to accurately estimate each individual cell type. Finally, although single-cell DNAm profile is a direct source of cell-type-specific methylation, its current utility for reference construction may be limited by the relatively poor data quality, including sparse genomic coverage, technical noise, and incomplete representation of relevant cell populations (5). In comparison, single-cell RNA-Seq technologies are more mature, with abundant tissue-specific cell atlases available as indirect reference resources.

To address these challenges, we developed HOMED (Hierarchically Optimized Methylation Deconvolution), a framework for DNAm deconvolution across a wide range of tissues and closely related cell types. HOMED first uses FACS-sorted DNAm cells to identify candidate CpG markers through hierarchical differential methylation analysis. Then, in the optimization step, bulk RNA-seq samples with paired DNAm data are iteratively deconvolved by a scRNA-seq-based deconvolution method. The resulting cell-type proportions serve as the pseudo-ground-truth to guide iterative optimization of CpG libraries on the paired bulk methylation samples. In particular, HOMED incorporates hierarchical relationships among cell types that share the same differentiation lineage, such that related cell types are first estimated as a group and subsequently resolved within each nested group. We benchmarked HOMED against other methods that utilize scRNA-seq or single-cell DNAm data as the guide and demonstrated significant improvement in cell-type proportion estimation across multiple tissue types. Finally, by applying HOMED to placental DNAm data from a smoking-exposed cohort, we identified smoking-associated differences in cell-type composition. Accounting for these compositional differences reduced inflated smoking-associated CpGs and improved the identification of biological pathways and functions associated with smoking, demonstrating the biological relevance and interpretability enabled by HOMED.

## Methods

### HOMED software, running environment and parameter tuning

HOMED was implemented in R and depends on the limma, lumi, nlme, genefilter, matrixStats, EpiDISH, doParallel, foreach, parallel, ggplot2, dplyr, and RColorBrewer packages. Reference construction was performed using HOMED_Reference(), which takes purified cell-type methylation profiles and annotations, training bulk methylation profiles, training cell-type proportions, and the predefined cell-type hierarchy as input. Deconvolution of independent bulk DNA methylation samples was subsequently performed using HOMED_Estimate() with the optimized hierarchical reference.

For reference construction, candidate CpGs were identified by differential methylation analysis at each hierarchical level using a significance threshold of *P* < 0.05, with up to 500 candidate probes selected per cell type. Candidate libraries were subsequently optimized against the training cell-type proportions using the iterative IDOL procedure. The maximum number of optimization iterations was set to 20, and optimization was terminated when the improvement in RMSE between successive iterations was <0.001. A final CpG library size of 500 probes was used. To improve computational efficiency during optimization, candidate probes were evaluated in batches; batch sizes ranging from 1 to 50 were assessed empirically, and a batch size of 5 was selected based on the trade-off between predictive performance and computational efficiency. All stochastic procedures were performed using a fixed random seed of 42 to ensure reproducibility. The HOMED source code, installation instructions, and a complete example demonstrating reference construction and application are publicly available through the HOMED GitHub repository.

### Benchmarking

For each real tissue dataset, pseudo-ground-truth cell-type proportions were first generated for all matched bulk RNA-seq and DNAm samples using DWLS/SCDC. The samples were then randomly divided into training and testing sets at a 7:3 ratio. CpG selection for HOMED was performed exclusively on the training set to prevent information leakage from the testing set.

For EpiSCORE, scRNA-seq counts were filtered to remove genes with no detectable expression, and then normalized to the median library size. Cell annotations were harmonized to the cell types evaluated in each tissue, and cell-type-specific expression references were constructed using ConstExpRef(). DNAm references were then imputed from the expression references using the SCM2 and RMAP databases with ImputeDNAmRef() and combined using ConstMergedDNAmRef(). Testing DNAm β-values were aggregated to promoter/TSS-level methylation using constAvBetaTSS(), and cell-type proportions were estimated using weighted robust partial correlation regression (wRPC; useW = TRUE, wth = 0.2, maxit = 1, 000). The same reference and parameters were used for simulated DNAm mixtures.

For scDeconv, scRNA-seq counts were converted to a Seurat object, normalized using NormalizeData(), and cell annotations were harmonized to the cell types evaluated in each tissue. Fifty pseudobulk profiles were generated using prepseudobulk(), and an scRNA-seq reference was constructed with scRef() using the pseudobulk profiles together with the matched bulk RNA-seq training data. The resulting RNA reference and training bulk RNA-seq data were then integrated with the corresponding training DNAm profiles using epDeconv(). Cell-type proportions in the held-out DNAm testing samples were estimated with 10 learners (learnernum = 10), and rescaling enabled (resscale = TRUE).

For scMD, single-cell DNAm data were used to construct tissue-specific methylation signatures. Single-cell clusters were mapped to the cell types evaluated in each tissue, and methylation β-values were calculated from methylated and unmethylated read counts. Cell-type-specific marker CpGs were identified using one-versus-rest Fisher’s exact tests, and the top 100 markers per cell type were selected using get_sig(). Bulk DNAm samples were then deconvolved using five methods implemented in scMD (CIBERSORT, EPIC, FARDEEP, DCQ, and Houseman), and their predictions were integrated using the scMD ensemble framework to obtain the final cell-type proportion estimates. The same signatures and parameters were applied to real and simulated DNAm samples.

### Evaluation metrics

The resulting references were applied to estimate cell-type proportions from the testing DNAm samples. Performance of all four methods was evaluated using Pearson correlation coefficient (PCC) and root mean squared error (RMSE) between estimated and ground-truth cell-type proportions for simulated datasets, or RNA-seq-derived pseudo-ground-truth proportions for real datasets. PCC and RMSE were calculated separately for each cell type and subsequently averaged to obtain dataset-level performance metrics. Negative PCC values were retained, whereas undefined PCC values resulting from constant predictions were excluded from mean PCC calculations. RMSE values were retained for all evaluated cell types, including those with undefined PCCs.

### Visualization of CpG markers generated by HOMED

To visualize the cell-type specificity of CpG markers selected by HOMED, we calculated the difference in methylation beta values (Δβ) between each target cell type and the mean of all other cell types at the same hierarchical level using the HOMED reference profiles. For each cell type, CpGs were ranked by the absolute magnitude of Δβ, and the top 50 CpGs were selected for visualization. For each selected CpG, methylation values were centered by subtracting the mean across cell types. The signs of centered values for hypomethylated markers were reversed so that positive values consistently represented target cell-type-specific methylation patterns. Markers were grouped by target cell type and methylation direction, with promoter-associated gene annotations included where available.

### Generation of simulated datasets

Simulated bulk DNAm datasets were generated from tissue-matched FACS-sorted methylation references. Note that the simulated datasets were used purely for testing and were not used in training references for each tissue. For each simulated sample, three purified methylation profiles were randomly selected per cell type. Cell-type proportions were sampled from a symmetric Dirichlet distribution, Dirichlet(1, …, 1) (8), and within each cell type the selected purified samples were assigned relative weights drawn from an additional symmetric Dirichlet distribution. The resulting sample-level weights were scaled to the corresponding cell-type proportions, and simulated bulk methylation β-values were generated as weighted linear combinations of the selected purified-cell profiles. A total of 100 simulated samples were generated per tissue, and the predefined cell-type proportions were retained as ground truth for performance evaluation.

### Real peripheral blood mononuclear cells (PBMCs) datasets

PBMC benchmarking used paired bulk DNAm and gene expression datasets from GSE105124 (9), GSE49065 (10) and GSE40736 (11). Processed methylation beta-values were directly downloaded and used for analysis, and raw microarray expression data were processed separately for each dataset, mapped to gene symbols using platform-specific annotations, and normalized using limma. When multiple probes mapped to the same gene, probe-level expression values were collapsed to gene-level values using the collapseRows method (12). Single-cell RNA-seq reference profiles were obtained from the 10x Genomics PBMC 68k dataset (13). FACS-sorted leukocyte methylation references were obtained from GSE167998 (6) and included 40 fluorescence-activated cell sorting-purified human leukocyte samples. For scMD, the scDNAm data for PBMCs were obtained from Zhou et al. (14). PBMC cell types were organized hierarchically, with monocytes, T cells, B cells, and NK cells at Level 1 and CD4⁺/CD8⁺ T cells at Level 2.

### Real lung datasets

Lung benchmarking used bulk DNAm and RNA-seq data from TCGA-LUAD (15). Raw RNA-seq counts were normalized using edgeR. Single-cell RNA-seq reference signatures were obtained from Sikkema et al. (16). FACS-sorted methylation references were compiled from four sources: 30 purified immune-cell methylomes from GSE35069 (17), 39 pulmonary endothelial-cell methylomes from GSE84395 (18), 3 epithelial-cell methylomes from GSE40699/ENCODE (19) and 14 normal fibroblast methylomes from GSE149958 (20). For scMD, the scDNAm data for lung tissue were also obtained from Zhou et al. (14). The resulting lung reference represented four major compartments for both Level 1 and Level 2 deconvolution: immune cells, epithelial cells, fibroblasts, and endothelial cells.

### Real placenta datasets

For the placenta benchmark, we utilized the bulk DNA methylation and gene expression datasets, the single-cell placenta atlas, and FACS-purified placental cell methylation data from the work with the cohort-level application on placenta research (21). The preprocessing and application pipeline were performed as reported.

### EWAS analysis of placentas exposed by maternal smoking

We analyzed the placental DNA methylation dataset GSE169598(22), including 72 smokers and 24 nonsmokers. Cell-type proportions were estimated using HOMED. Smoking-associated CpGs were identified with limma before and after adjustment for estimated cell-type proportions, together with clinical variables including gestational age and fetal sex. KEGG pathway enrichment was performed using missMethyl(23) to compare biological signals before and after cell-type adjustment.

## Results

### Overview of HOMED and benchmarking against other methods

HOMED is a hierarchical optimization framework for DNAm deconvolution. The sketch of the HOMED framework is shown in **Figure 1a**. It estimates cell-type composition in a coarse-to-fine manner: broad parent cell types that share a common differential lineage are first resolved, followed by finer cell types or subtypes within each lineage. HOMED then iteratively refines the estimation of each lineage, cell type, or subtype by prioritizing CpG probes that improve estimation accuracy in heterogeneous mixtures. In each iteration, the selected CpG probe set is optimized and refined, guided by the pseudo-ground-truth of the training samples. Specifically, these pseudo-ground-truth proportions are derived from applying state-of-the-art scRNA-seq reference-based deconvolution methods to paired bulk gene expression data. To ensure biological consistency, subtype proportions are rescaled so that their proportions sum to the corresponding parent cell-type proportion. By incorporating biological cell-type relationships into both deconvolution and probe optimization, HOMED aims to improve the resolution of closely related cell types or subtypes with highly similar DNAm signatures.

**Figure 1.**
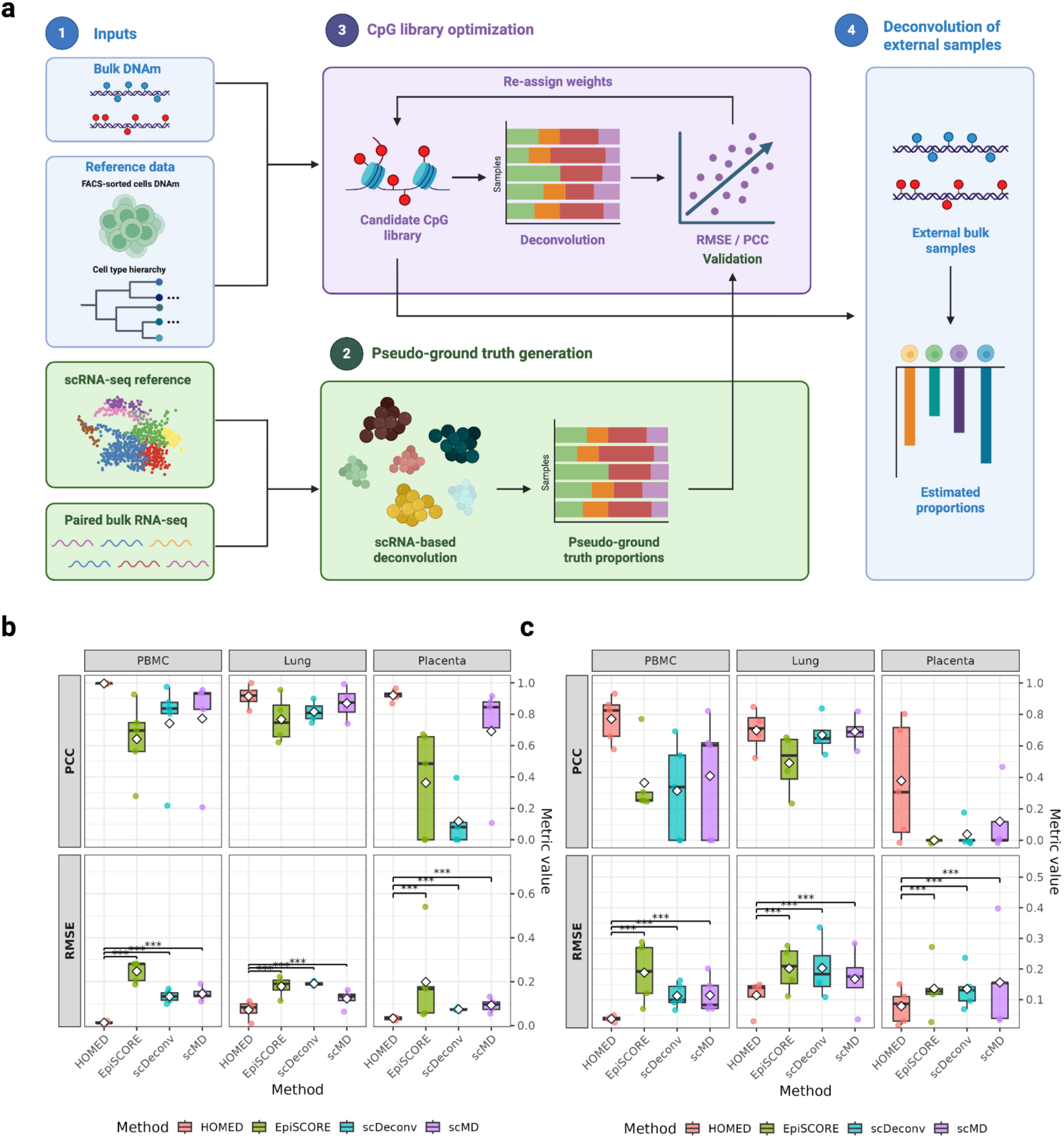
Illustration of the HOMED framework and its benchmark over other deconvolution methods. (a) Sketch of the HOMED framework. (1) Inputs: FACS-sorted cell DNAm profiles and cell-type hierarchy are combined with paired bulk DNAm, while matched scRNA-seq reference data and paired bulk RNA-seq samples are used for pseudo-ground-truth generation. (2) Pseudo-ground-truth generation: bulk RNA-seq samples are deconvolved by DWLS or SCDC using the scRNA-seq reference to estimate cell-type proportions, which serve as pseudo-ground-truth labels for optimization. (3) CpG library optimization: candidate cell-type-discriminatory CpGs are selected from FACS-sorted DNAm references and iteratively evaluated through deconvolution of training bulk DNAm samples. CpG selection probabilities are updated based on deconvolution performance, measured by RMSE and PCC relative to the pseudo-ground-truth proportions, resulting in an optimized CpG library. (4) Deconvolution of external samples: the optimized DNAm reference library is applied to independent bulk DNAm datasets to estimate cell-type proportions. (b) Boxplots to show overall performance comparison of HOMED, EpiSCORE, scDeconv, and scMD on simulated PBMC, placenta, and lung tissues. Each point represents a cell type in the tissue, and diamonds indicate mean values across cell types in that tissue. Metrics shown are PCC and RMSE. (c) Boxplots to show overall performance comparison of HOMED, EpiSCORE, scDeconv, and scMD on real PBMC, placenta, and lung tissues. Each point represents a cell type in the tissue, and diamonds indicate mean values across cell types in that tissue. Metrics shown are PCC and RMSE.

We benchmarked HOMED against three single-cell-guided DNAm deconvolution methods: EpiSCORE and scDeconv, which use scRNA-Seq data as the guide similar to HOMED, and scMD, a more recent method using scDNAm data as the guide. Earlier methods that do not use single-cell data were not included to maintain a fair comparison with state-of-the-art single-cell-guided approaches. We evaluated all methods on both simulated mixtures and real DNAm datasets from three tissue types: PBMC, placenta, and lung tissues. For the real datasets, pseudo-ground-truth cell-type proportions were derived by deconvolving bulk gene expression data using DWLS (24) or SCDC (25) from the same samples with paired DNAm measurements.

Across all datasets, HOMED consistently achieved the best accuracy, demonstrated by the lowest root mean square error (RMSE) and highest Pearson correlation coefficient between the estimated and scRNA-seq-derived pseudo-ground truth **(Figure 1b**, **1c, Supplementary Table 1)**. Specifically, in the simulated datasets, HOMED achieved the lowest mean RMSEs of 0.01, 0.07, and 0.03 for PBMC, lung, and placenta respectively, whereas the other three methods’ RMSEs were 1.7 to 25 times those of HOMED, with EpiSCORE yielding the worst RMSEs in general. In these same datasets, HOMED yielded the highest mean PCCs of 1.00, 0.92, and 0.93 for PBMC, lung, and placenta respectively, compared to the other three methods with much worse PCCs ranging from 0.17 to 0.87. In particular, scDeconv showed the lowest PCC of 0.17 in the simulated placenta data. On the corresponding real datasets, all methods’ performance decreased compared to the simulated data, as expected because real-world samples contain greater biological and technical variability than controlled simulated mixtures. Nevertheless, HOMED maintained superior performance, achieving the lowest mean RMSEs of 0.04, 0.12, and 0.06, and the highest PCCs of 0.77, 0.69, and 0.41, for PBMC, lung, and placenta tissues respectively. In comparison, the other three methods have RMSE values 1.42 to 4.75 times those of HOMED, ranging from 0.11 to 0.20, and dramatically worse PCCs ranging from -0.26 to 0.69. Notably, all three competing methods yielded negative mean PCCs for real placenta data, whereas HOMED was the only method to achieve a positive mean PCC of 0.41.

### HOMED performs the best deconvolution in PBMC DNAm data

We further examined detailed cell-type-level performance across the four methods to determine how effectively HOMED resolves biologically related cell populations in both simulated and real PBMC datasets (**Figure 2**). In simulated PBMCs (**Figure 2a**), HOMED accurately recovers all immune cell types, including the closely related CD4+ and CD8+ T-cell subtypes, with PCCs of 0.99 to 1 and RMSEs of 0.01 to 0.02 across all of them. By contrast, EpiSCORE performs much worse and fails to estimate the variations mostly, with PCCs ranging from 0.28 to 0.93 and RMSEs from 0.19 to 0.28. It is especially problematic for estimating monocytes and CD8+ T cells. scDeconv, while showing decent trends as demonstrated by the PCCs of 0.22 to 0.97, has large prediction errors shown by the RMSEs of 0.1 to 0.17. scMD also shows inferior performance with PCCs of 0.21 to 0.95 and RMSEs of 0.11 to 0.19. All three methods, EpiScore, scDeconv, and scMD struggle in estimating CD8+ T cell proportions, which HOMED does superbly well.

**Figure 2.**
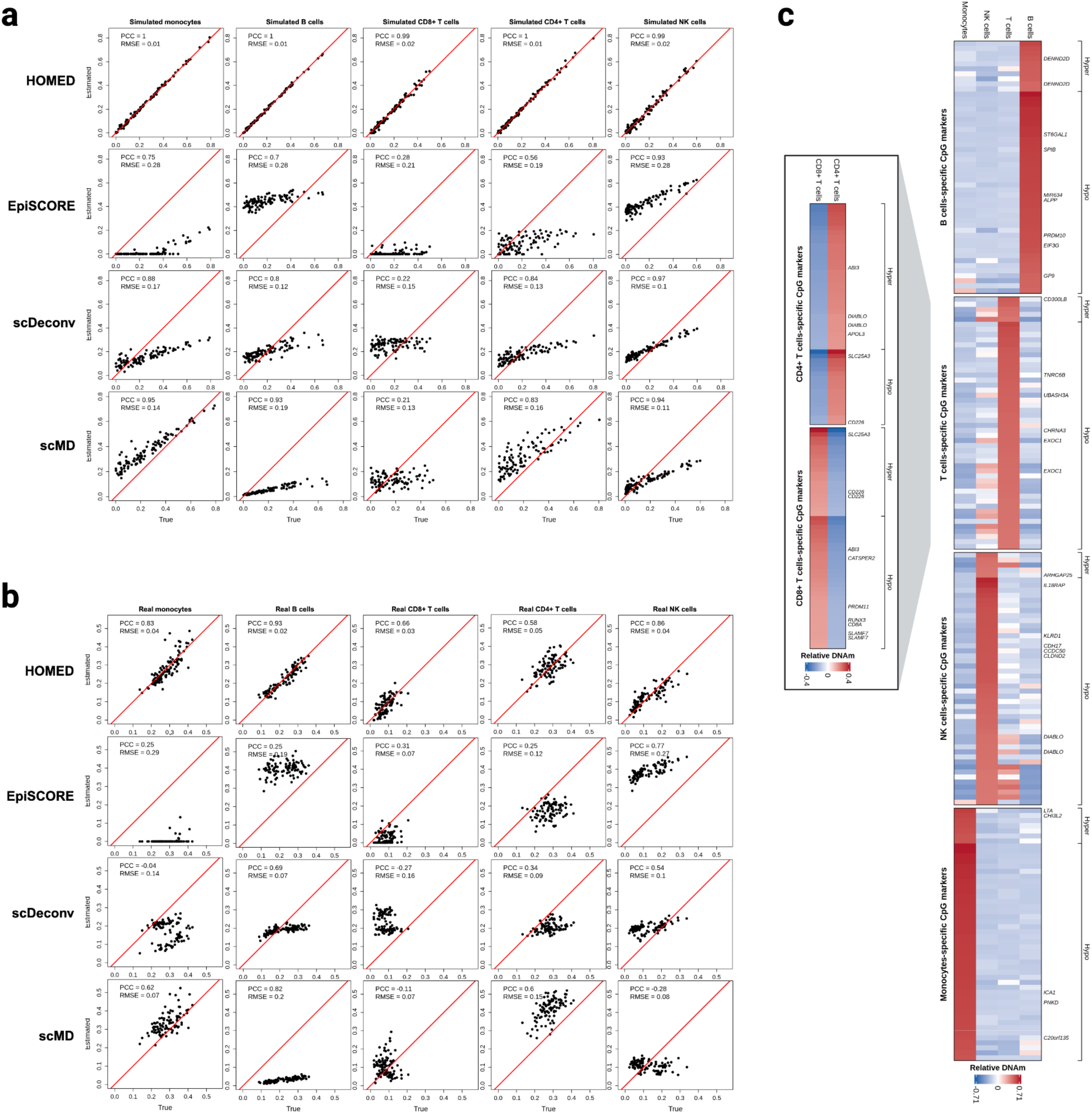
Comparison of cell-type level performance of four deconvolution methods on simulated and real PBMC tissues. HOMED was benchmarked against EpiSCORE, scDeconv, and scMD, three different state-of-the-art single-cell-guided DNAm deconvolution methods that use either scRNA-Seq or scDNAm data as the guide. (a) Scatter plots comparing estimated (y-axis) and true (x-axis) cell-type proportions for five major PBMC cell types (B cells, CD4+ T cells, CD8+ T cells, monocytes, and NK cells) in simulated PBMC samples. Results are shown for HOMED, EpiSCORE, scDeconv, and scMD. (b) Corresponding results for real PBMC samples, with cell-type proportions derived from paired bulk gene expression data used as pseudo-ground truth. The red diagonal line indicates perfect agreement between estimated and true or pseudo-ground-truth proportions (y = x). PCC and RMSE are reported in each panel to quantify deconvolution performance. (c) Heatmaps of HOMED-selected cell-type-specific CpG markers. The left heatmap shows markers distinguishing CD4+ and CD8+ T-cell subtypes, and the right heatmap shows markers for the four major PBMC lineages (B cells, T cells, NK cells, and monocytes). For visualization, methylation beta values are normalized across cell types for each CpG, and presented as mean-centered, direction-normalized methylation beta values. Positive (red) values indicate persistent methylation pattern, characteristic of biomarkers of the interested cell type, whereas negative (blue) values indicate the opposite pattern.

In real PBMC data, HOMED continues to perform the best, even though all methods have decreased performance as compared to the simulated data (**Figure 2b**). HOMED shows PCCs in the range of 0.58 to 0.93 and RMSEs of 0.02 to 0.05. Even in the two similar CD4+ and CD8+ cell subtypes, HOMED achieves very decent performance, with PCC/RMSE of 0.66/0.03 for CD8+ T cells and 0.58/0.05 for CD4+ T cells. In comparison, EpiScore, scDeconv and scMD perform much worse than HOMED. EpiSCORE shows PCCs in the range of 0.25 to 0.77, and RMSE of 0.07 to 0.29; it fails to estimate monocytes, systematically overestimates B cells and NK cells, and underestimates CD4+ T cells. scDeconv presents PCCs in the range of -0.27 to 0.69, and an RMSE of 0.07 to 0.16. Strangely, the estimation of monocytes and CD8+ T cells shows split trends, rather than expected monotonic trends. scMD presents PCCs in the range of -0.28 to 0.82, and an RMSE of 0.07 to 0.20. It appears to really struggle with CD8+ T cells and NK cells, both having negative PCCs. Moreover, it consistently underestimates B cells but overestimates CD4+ T cells. Similar to the simulated data, all three methods, EpiScore, scDeconv and scMD have a lot of trouble estimating CD8+ T cell proportions.

To further examine the cell-type-specific methylation signatures underlying HOMED’s deconvolution performance, we visualized the CpG markers selected by HOMED across the PBMC hierarchy **(Figure 2c)**. For PBMCs, at the cell type level, CpG markers selected for B cells, T cells, NK cells, and monocytes show distinct cell-type-specific methylation patterns. Importantly, this pattern is also observed at the cell subtype level, separating the closely related CD4+ and CD8+ T-cell subtypes. These results directly illustrate the success of the hierarchical strategy used by HOMED to identify informative CpG markers for both the cell types and their subtypes.

### HOMED performs the best deconvolution in lung DNAm data

Lung tissues are heterogeneous and comparatively less studied compared to PBMC, thus benchmarking on lung tissue provides additional support for tissue generalizability of HOMED. We followed the annotation scheme of EpiSCORE and categorized the cell types into epithelial, endothelial, stromal, and immune cells (3). In simulated lung tissues (**Figure 3a)**, HOMED has the most accurate estimation of each cell type, reaching an average PCC of 0.92 (range 0.82-1.00) and an average RMSE of 0.07 (range 0.01-0.11). EpiSCORE, on the contrary, has the most dispersed and inaccurate estimation among the four methods. It yields average PCC of 0.77 (range 0.62-0.95) and RMSE of 0.18 (range 0.11-0.22); scDeconv reaches an average PCC of 0.82 (range 0.75-0.90) and RMSE of 0.19 (range 0.18-0.20); and scMD generates the second best results, with an average PCC of 0.87 (range 0.74-0.99) and RMSE of 0.12 (range 0.06-0.16).

**Figure 3.**
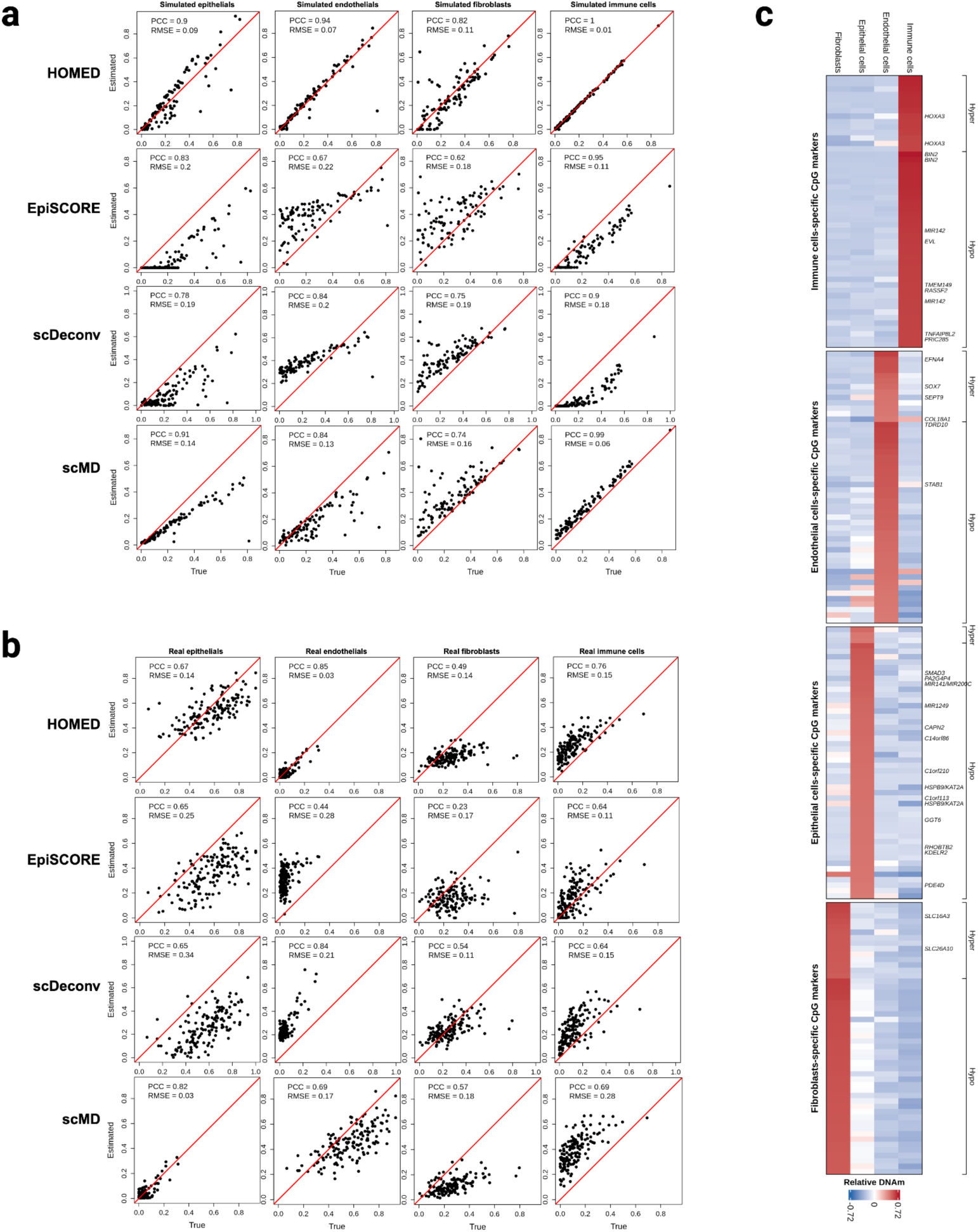
Comparison of cell-type level performance of four deconvolution methods on simulated and real lung tissues. HOMED was benchmarked against EpiSCORE, scDeconv, and scMD. **(a)** Scatter plots comparing estimated (y-axis) and true (x-axis) cell-type proportions for four major lung cell populations (epithelial cells, endothelial cells, fibroblasts, and immune cells) in simulated lung samples. **(b)** Corresponding results for real lung samples, with cell-type proportions derived from paired bulk gene expression data used as pseudo-ground truth. The red diagonal line indicates perfect agreement between estimated and true or pseudo-ground-truth proportions (y = x). PCC and RMSE are reported in each panel to quantify deconvolution performance. **(c)** Heatmap of HOMED-selected cell-type-specific CpG markers for the four major lung cell populations. The coloring scheme is the same as Figure 2.

The performance of all methods in the real lung tissue data decreased as expected. However, HOMED maintains the best method overall (**Figure 3b, Supplementary Table 1**). It reaches an average PCC of 0.69 (range 0.49-0.85) and an average RMSE of 0.12 (range 0.03-0.15). EpiSCORE again has the worst performance among the four methods, yielding an average PCC of 0.49 (range 0.23-0.65) and RMSE of 0.20 (range 0.11-0.28). It has the most difficulty in estimating endothelial cells. scDeconv reaches an average PCC of 0.67 (range 0.54-0.84) and RMSE of 0.20 (range 0.11-0.34); scMD generates the second best results, with an average PCC of 0.69 (range 0.57-0.82) and RMSE of 0.17 (range 0.03-0.28) Together, these results consistently show that HOMED has the most robust estimation on all cell types in lung samples. Additionally, the cell-type-specific CpG markers selected by HOMED show the distinct methylation patterns across the four major lung cell populations (**Figure 3c**), confirming its robust deconvolution performance in both simulated and real lung tissues.

### HOMED performs the best deconvolution in the placental tissue

The placenta is a very heterogeneous and blood-rich organ that is present transiently during pregnancy. It is composed of a trophoblast surface and villous core (26). Trophoblast cells cover the placenta villus, the basic units of placentas. Major trophoblast populations include syncytiotrophoblasts, villous cytotrophoblasts, and extravillous trophoblasts, which are highly related and difficult to deconvolve (21, 27). The villous core contains Hofbauer cells, stromal fibroblasts, endothelial cells and the nuclei-present red blood cells (nRBCs) from the baby within the endothelium. In short, the human placenta presents some of the most complex human tissues, and deconvolution is thus expected to be very challenging.

In the placenta data, HOMED consistently outperforms the other three methods across a diverse set of placental cell populations, including trophoblast subtypes (extravillous trophoblasts, villous trophoblasts, and syncytiotrophoblasts), as well as stromal, endothelial, Hofbauer, and nRBCs (**Figure 4**). In particular, across all cell types in simulated placenta data, HOMED achieves an impressive overall PCC of 0.93 (range 0.86-0.99) and RMSE of 0.03 (range 0.01-0.04) and closely recovers the ground truth, as shown in **Figure 4a**. In comparison, EpiSCORE performs worse, with a much lower average PCC of 0.60 (range 0.49-0.67), calculated across the five cell types with defined PCC values, and average RMSE of 0.18 (range 0.05-0.54), across all seven cell types. Notably, EpiSCORE fails to estimate endothelial and syncytiotrophoblast proportions, assigning zero proportions to both cell types across all samples, resulting in undefined PCC values that were excluded from the mean PCC calculation. Additionally, EpiSCORE substantially overestimated villous trophoblast proportions. scDeconv has the worst performance, with an average PCC of 0.17 (range -0.17-0.95) and RMSE 0.07 (0.07-0.08), and significantly fails to recover the variance of all cell types. scMD again is the second best method, achieving an average PCC of 0.69 (range 0.11-0.92) across the five cell types with valid PCC values, and average RMSE of 0.09 (0.05-0.13) across all cell types. Most importantly, scMD does not have extravillous trophoblasts nor nRBCs in the original scDNAm atlas (16), thus can not estimate these two cell types. Indeed, lacking certain reference cell types is a major technical issue of applying scMD currently.

**Figure 4.**
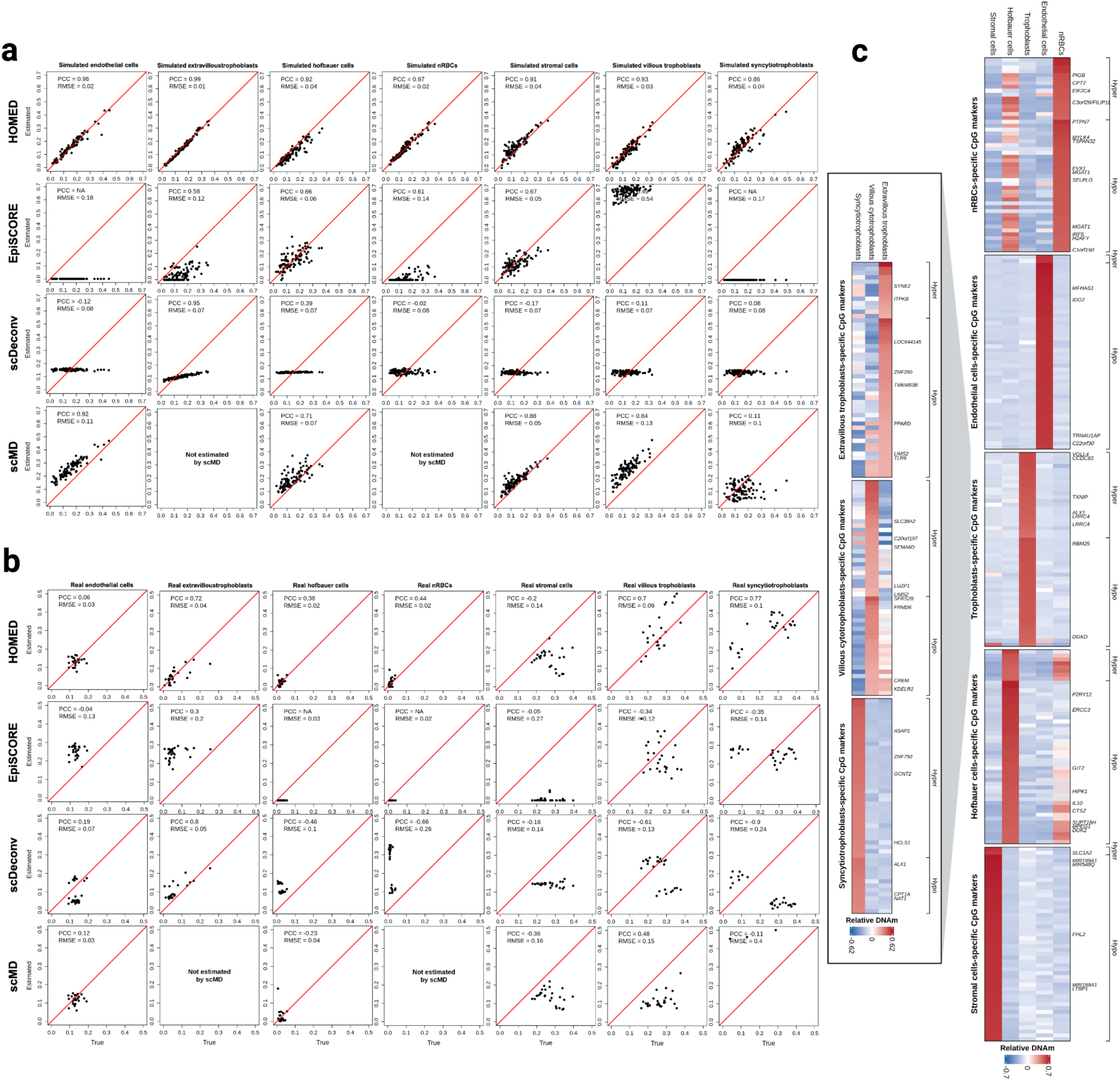
Comparison of cell-type level performance of four deconvolution methods on simulated and real placenta tissues. HOMED was benchmarked against EpiSCORE, scDeconv, and scMD. **(a)** Scatter plots comparing estimated (y-axis) and true (x-axis) cell-type proportions for seven placenta cell populations (endothelial cells, Hofbauer cells, nucleated red blood cells [nRBCs], stromal cells, extravillous trophoblasts, villous cytotrophoblasts, and syncytiotrophoblasts) in simulated placenta samples. **(b)** Corresponding results for real placenta samples, with cell-type proportions derived from paired bulk gene expression data used as pseudo-ground truth. The red diagonal line indicates perfect agreement between estimated and true or pseudo-ground-truth proportions (y = x). PCC and RMSE are reported in each panel to quantify deconvolution performance. **(c)** Heatmaps of HOMED-selected cell-type-specific CpG markers at the two levels of the placenta hierarchy. The heatmaps show markers distinguishing the major placenta cell lineages and markers distinguishing the three trophoblast subtypes. The coloring scheme is the same as in Figure 2.

We next compared the four methods on a combined placenta dataset with paired DNA methylation and transcriptomics data (28, 29), where the ground truth of cell proportions is inferred from the transcriptomics data modality (**Figure 4b**). As expected, the performance of these methods all decreases in the more complex real data scenario. Still, HOMED achieves the best accuracy across the major placental populations, with an average PCC of 0.41 (range: -0.20-0.77) and an average RMSE of 0.06 (range: 0.02-0.14) across all seven major cell types. Worth noticing, HOMED deconvolves the three types of trophoblast cells very well, achieving cell-type-specific PCCs of 0.72, 0.70 and 0.77 for extravillous trophoblasts, villous trophoblasts, and syncytiotrophoblasts, respectively. In comparison, EpiSCORE shows much poorer performance, achieving an average PCC of -0.10 (range: -0.35-0.30), calculated across the five cell types with valid PCC values, and an average RMSE of 0.13 (range: 0.02-0.27). In particular, it estimates Hofbauer cells and nRBC proportions to be all zero; thus, their PCCs are shown as NAs, reflecting incomplete representation of placental cell types in its atlas framework. scDeconv produces an average PCC of -0.26 (range: -0.90-0.80) and RMSE of 0.14 (range: 0.05-0.26). It again shows the split patterns of estimation for most cell types, instead of the expected monotonic pattern. scMD achieves the second-highest average PCC of -0.02 (range: -0.36-0.48), calculated across the five cell types with valid PCC values, but the highest mean RMSE of 0.16 (range: 0.04-0.40). Similar to the simulated placentas, scMD cannot estimate extravillous trophoblasts or nRBCs due to missing these cell types in the scDNAm atlas (16). In conclusion, despite the significant challenge due to tissue complexity such as placentas, HOMED is still able to perform decently and does particularly well in the three specialized trophoblast subtypes, which are pivotal for placental biology.

The HOMED-selected CpG markers also reveal unique methylation patterns across the placenta hierarchy (**Figure 4c**). At the cell type level, cell-type specific CpG signatures clearly separate the major placental populations, including nRBCs, endothelial cells, trophoblasts, Hofbauer cells, and stromal cells. Within the trophoblast lineage, a subset of markers further distinguish the highly similar extravillous trophoblasts, villous cytotrophoblasts, and syncytiotrophoblasts, confirming its advantage in resolving these trophoblast subtypes despite their shared lineage.

### HOMED improves the epigenome-wide association study of the effect of smoking on placentas

We further demonstrate the practical value of DNA methylation deconvolution by HOMED by examining its impact on downstream EWAS (epigenome-wide association analysis). Towards this end, we use an Illumina EPIC DNA methylation dataset from placentas of 72 smokers and 24 nonsmoker controls (22) (**Figure 5**). As a proof of principle, cell-type proportions directly annotated from an independent single-nucleus RNA-seq dataset of 32 placentas show strong concordance with HOMED-estimated proportions in the methylation cohort (PCC = 0.87, P < 0.05; **Figure 5a**). HOMED further identifies significant smoking-associated differences in endothelial, nucleated red blood cell (nRBC), stromal, and villous trophoblast proportions (P < 0.05; **Figure 5b**), confirming that maternal smoking is accompanied by substantial remodeling of placental cellular composition.

**Figure 5.**
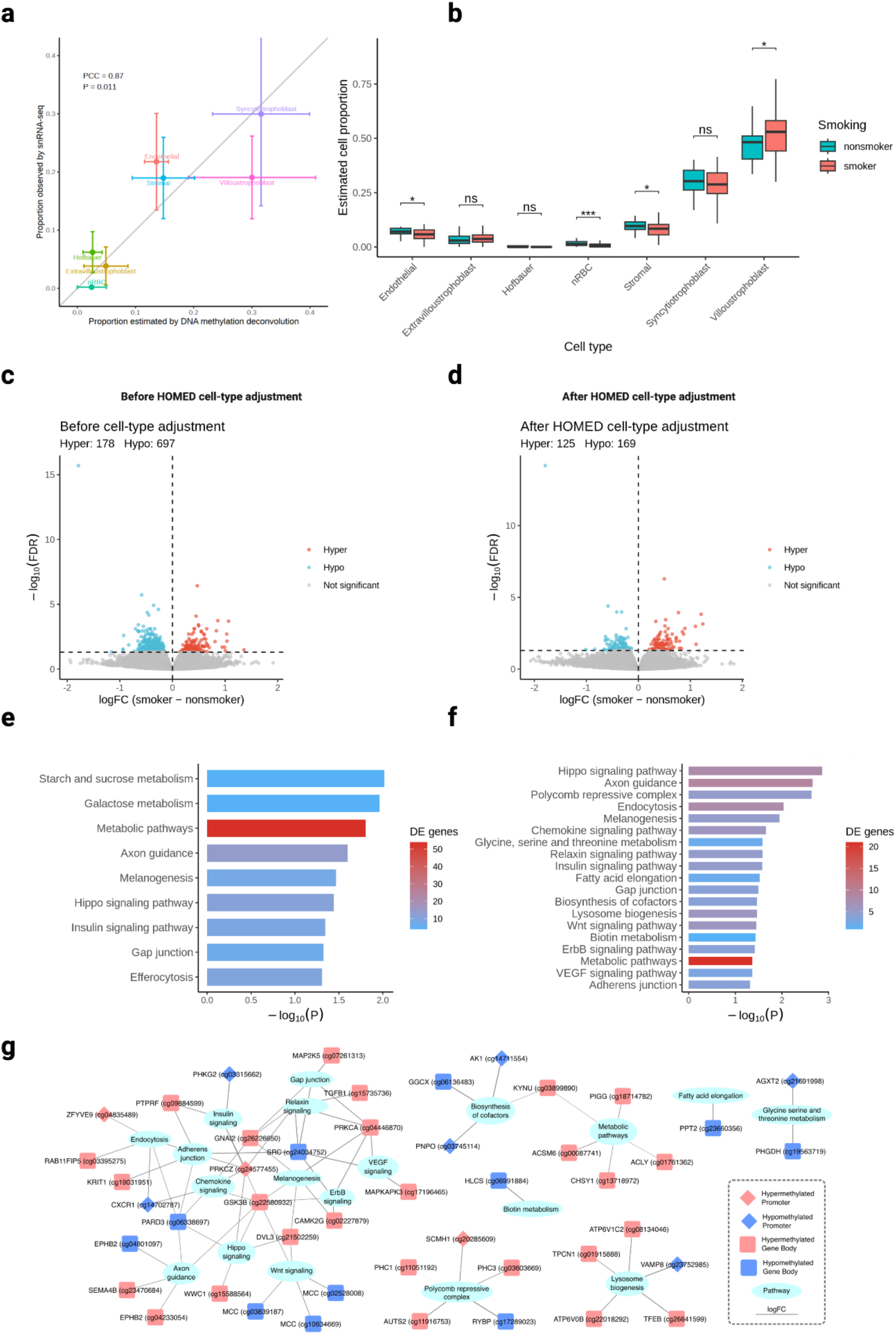
HOMED enhances detection of smoking-associated epigenetic signals. (a) Comparison of cell-type proportions estimated by DNA methylation deconvolution and directly annotated by snRNA-seq data. Points indicate mean proportions; horizontal and vertical error bars show standard deviation in methylation-based placental cell estimates and snRNA-seq cell proportions, respectively. The dotted line indicates *y* = *x*, with PCC and *P* value shown. (b) HOMED-estimated cell-type proportions in smoker and nonsmoker placentas; The significance level is indicated above each comparison (n.s., not significant; *P < 0.05; ***P < 0.001). (c-d) Volcano plots of smoking-associated CpGs before (c) and after (d) cell-type adjustment, showing hypermethylated, hypomethylated, and nonsignificant CpGs. (e-f) KEGG pathway enrichment of smoking-associated CpGs before (c) and after (d) HOMED cell-type adjustment; bar length represents -log10(*P*) and color indicates the number of associated CpGs. (g) Network of enriched pathways and their top 5 smoking-associated CpGs after cell-type adjustment by logFC. Node color indicates hyper- or hypomethylation, node shape indicates genomic region class, and logFC-weighted edges connect CpGs to associated pathways.

The impact of cell-type adjustment is further shown in the downstream EWAS analysis. Without adjustment for these compositional differences, EWAS identifies 875 significant CpGs after controlling for gestational age and fetal sex (**Figure 5c**). Once incorporating HOMED-estimated cell proportions, the number of significant CpGs decreases by approximately three-fold to 294 CpGs (**Figure 5d**). This reveals that a substantial fraction of previous smoking-associated methylation signals in **Figure 5c** reflects underlying cell-type heterogeneity rather than the intended CpG level difference. Pathway over-representation analysis further shows a marked shift in biological interpretation (**Figure 5e, f**). Before cell-type adjustment, only 9 nominally enriched pathways remained after exclusion of disease-specific terms, including broad carbohydrate and metabolic processes such as galactose metabolism, starch and sucrose metabolism, and general metabolic pathways (**Figure 5e**). In contrast, after HOMED adjustment, as many as 19 nominally enriched pathways emerged, including Polycomb repressive complex, Wnt, Hippo, VEGF, ErbB, adherens junction, and axon guidance signaling (**Figure 5f**). These pathways collectively reveal developmental epigenetic regulation, trophoblast differentiation and migration, and placental angiogenesis under the impact of maternal smoking. These pathways converge on smoking-associated methylation changes in key regulators (**Figure 5g**). This includes GSK3B (cg22580932) linking Wnt/Hippo signaling, PRKCA (cg04446870) and SRC (cg24034752) connecting VEGF/ErbB signaling, and Polycomb components such as PHC1 (cg11051192) and SCMH1 (cg20285609) involved in developmental chromatin regulation.

Additional examples of the major value of HOMED on EWAS are also demonstrated in another study of preeclamptic placentas (21). It shows improved confounding adjustment, drastically reducing inflation of differential methylation in heterogeneous bulk tissues, and revealing disease-associated methylation signals which would otherwise be obscure.

### Computational efficiency and fine-tuning of HOMED

Computational scalability during CpG library optimization is an important consideration for HOMED, as candidate CpGs are iteratively evaluated to identify reference signatures that maximize deconvolution accuracy. To improve computational efficiency, HOMED incorporates batch-based CpG evaluation, in which multiple candidate CpGs are evaluated simultaneously during each iteration. We therefore assessed the trade-off between computational efficiency and deconvolution performance by varying the batch size from 1 to 50 across PBMC, lung, and placenta datasets (**Figure 6**). Across all three tissues, increasing the batch size substantially reduced optimization runtime, with the greatest improvement occurring at smaller batch sizes. This trend is expected because larger batches allow more candidate CpGs to be evaluated and incorporated during each optimization step, thereby reducing the total number of iterative evaluations required. The reduction in runtime gradually plateaued at larger batch sizes, indicating diminishing computational benefits from further increasing the number of CpGs evaluated per iteration.

**Figure 6.**
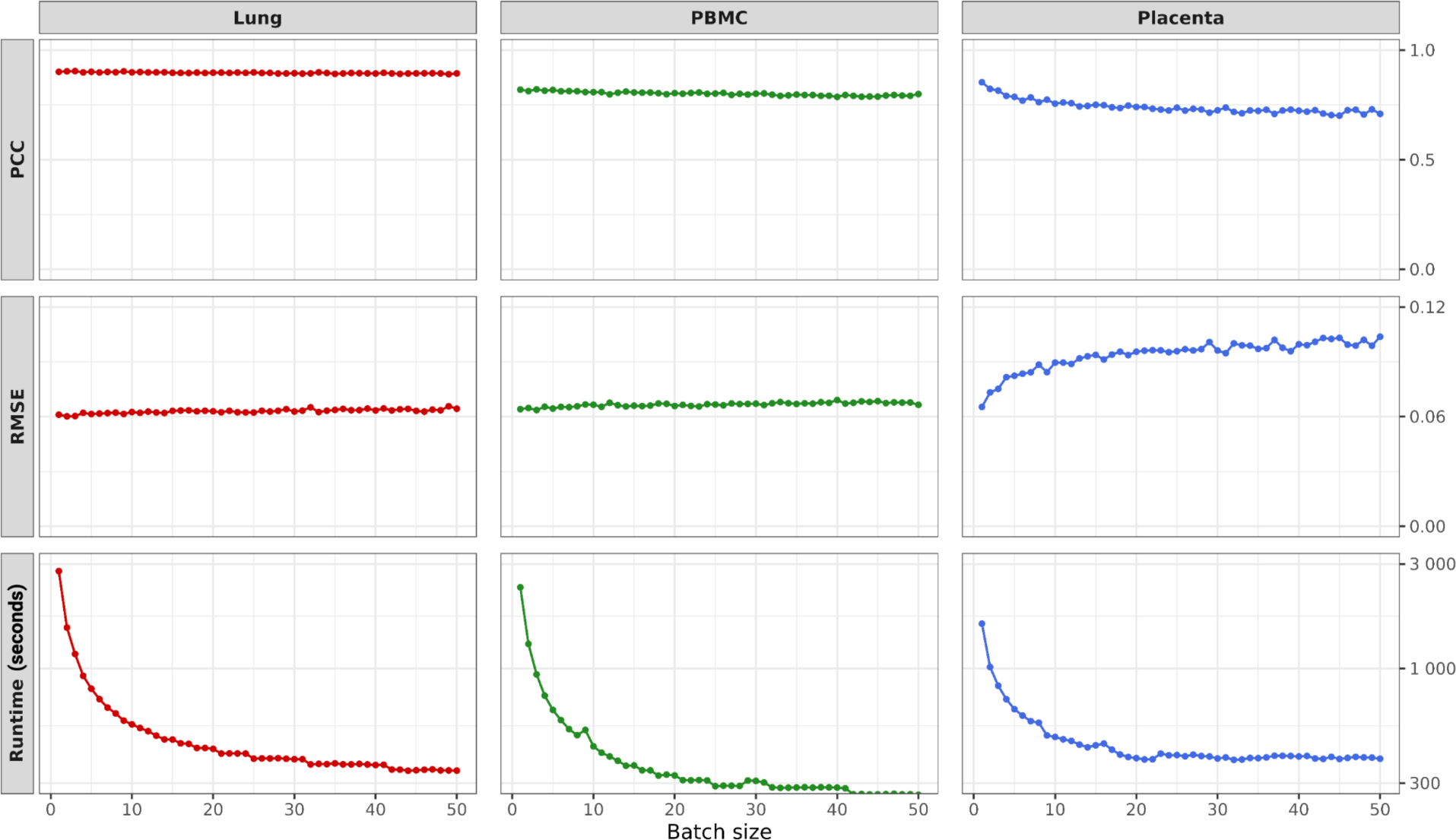
Effect of batch size on HOMED optimization performance. Average PCC, RMSE, and runtime were evaluated across batch sizes from 1 to 50. PCC and RMSE remained largely unchanged, while runtime decreased substantially with increasing batch size. A batch size of 5 was selected as the default for analyses to balance computational efficiency and deconvolution accuracy.

Overall, these results highlight a trade-off between computational efficiency and the granularity of CpG selection when determining batch size. Smaller batch sizes enable finer-grained optimization but incur greater computational costs, whereas larger batch sizes offer diminishing runtime improvements and can compromise accuracy, particularly for placenta. We therefore selected a batch size of 5 as the default setting in the HOMED package and used this value for all analyses reported in this study. This setting captures much of the computational speedup afforded by batching while maintaining deconvolution performance close to that achieved with smaller batch sizes across all three tissues.

## Discussion

Here we present HOMED, a hierarchical and multimodal optimization-based DNA methylation deconvolution method. Across simulated and real datasets from PBMC, lung and placenta, HOMED consistently achieved predominantly superior deconvolution accuracies, with particularly strong gains for closely related or poorly resolved cell populations. Rather than representing an incremental improvement in reference-based deconvolution approaches, HOMED introduces a different strategy for constructing methylation references: it combines biological cell-type hierarchy with iterative optimization guided by orthogonal transcriptomic information. This design allows CpG signatures to be selected not only for their ability to distinguish purified cell types, but also for their ability to recover cellular proportions in heterogeneous bulk tissues. This coarse-to-fine strategy is especially advantageous in tissues containing developmentally related cell types, such as CD4+ T and CD8+ T cells in PBMC and the various trophoblast cell types in placenta.

Among the three competing methods, scMD generally showed the second-best performance. Unlike EpiSCORE and scDeconv, which use scRNA-seq information to infer or guide methylation references, scMD directly leverages scDNAm data to construct cell-type-specific methylation signatures. This provides a more direct representation of cell-type-specific methylation patterns and may contribute to its relatively strong performance. However, this advantage is highly dependent on the availability of scDNAm reference atlas. In our placenta analyses, for example, the scDNAm reference used by scMD did not contain extravillous trophoblasts or nRBCs, preventing scMD from estimating these cell populations. More broadly, current scDNAm datasets remain substantially less abundant than scRNA-seq atlases, particularly for complex tissues and specialized cell populations. Thus, while scMD has the potential for DNAm deconvolution straightforwardly, its broad success will depend on making more comprehensive scDNAm reference atlases available, as well as improving their data quality. In contrast, HOMED leverages both the wide availability and higher quality of scRNA-seq data, as well as explicitly modeling the hierarchical resolution of closely related cell populations, contributing to its unanimous best performance in all tissues and conditions tested here.

Importantly, the value of improved deconvolution extends beyond estimation of cell proportions and directly influences the accurate selection of epigenetic biomarkers. In the example of the smoking-exposed placenta cohort, HOMED identified significant differences in placental cell type compositions between smokers and nonsmokers. Moreover, adjustment for these estimated proportions substantially altered the EWAS landscape. Cell-type adjustment reduced the inflation of significant CpGs while revealing more precisely the pathway-level signals involving placental development. Important biological functions related to smoking, including Wnt signaling, VEGF-mediated angiogenesis, and Polycomb-associated epigenetic regulation, are masked if the HOMED-based cell-type adjustment is not applied. These results illustrate how inaccurate or incomplete modeling of tissue composition can have a profound adverse impact on downstream biomarker studies by both inflating differential methylation signals and obscuring internal biological patterns. For examples, *GSK3B* (cg22580932), linking Wnt/Hippo signaling, is consistent with smoking-induced suppression of placental WNT signaling and trophoblast proliferation (30), *PRKCA* (cg04446870) and *SRC* (cg24034752), connecting VEGF/ErbB signaling, had also been implicated in placental methylation responses to maternal smoking (31), and Polycomb components such as PHC1 (cg11051192) and SCMH1 (cg20285609) involved in developmental chromatin regulation, were reported to be hypermethylated with smoking (32). Notably, EPHB2 (cg04233054), involved in axon guidance and migration networks, was also selected in all three placental smoking biomarker models in the original ECHO study (22), providing independent and strong support for the biological relevance of the HOMED-refined EWAS analysis.

Despite the major advantages of HOMED shown here, some future improvements remain. First, it currently focuses primarily on two-level hierarchical structures and may require additional validation to support more complex cellular ontologies and developmental trajectories. Second, pseudo-ground-truth proportions generated through RNA-based deconvolution are themselves subject to estimation uncertainty and may propagate bias into DNAm reference optimization; however, the large quantities of paired gene expression and DNAm samples should mitigate this. Third, the success of HOMED relies on the bridge of samples that have paired DNA methylation and gene expression measurements, in which the RNA-Seq data modality is used as ground truth to guide the DNA methylation deconvolution process. Moreover, in this study, the DNA methylation reference data are obtained from purified cell-type references, which may be mitigated with the improved quality of single-cell methylation references in the future.

In summary, HOMED is a next-generation, high-performing computational framework for integrating cellular hierarchy and cross-modal information into DNA methylation deconvolution. By improving both the resolution of closely related cell populations and the robustness of downstream epigenomic analyses, HOMED provides a practical yet powerful strategy for extracting cell-type-aware biological information from heterogeneous bulk DNA methylation data.

## Abbreviations

CpG: Cytosine-phosphate-guanine
DNAm: DNA methylation
FACS: Fluorescence-activated cell sorting
GEO: Gene Expression Omnibus
HOMED: Hierarchically Optimized Methylation Deconvolution
LUAD: Lung adenocarcinoma
MCA1: Mouse Cell Atlas-1
nRBC: Nucleated red blood cell
ORA: Over-Representation Analysis
PBMC: Peripheral blood mononuclear cell
PCC: Pearson correlation coefficient
RMSE: Root mean squared error
RNA-seq: RNA sequencing
SCDC: Single-cell Deconvolutional Consensus
scRNA-seq: Single-cell RNA sequencing
TCGA: The Cancer Genome Atlas.

## Declarations

### Data Availability

All datasets used in this study are publicly available through GEO and TCGA with accession numbers listed in the Methods section. HOMED is available for installation on GitHub (https://github.com/lanagarmire/HOMED). The source code, analyses, documentation, and reproducible benchmarking workflows are archived on Zenodo (https://doi.org/10.5281/zenodo.23487321).

## Acknowledgements

The authors thank members of the Garmire lab for helpful discussions and technical support during this work.

## Authors’ contributions

LG envisioned this project, obtained the funding, supervised the study, and revised the manuscript. YL obtained data, improved HOMED runtime, performed benchmarking, generated the figures, and wrote the initial manuscript. YD provided initial data, designed the benchmarking workflow, helped with the analysis, and revised the manuscript. YC collected paired bulk DNA methylation and gene expression data and reproduced the HOMED vignette.

## Supplementary Figures and Table Legends

**Supplementary Table 1.** Comparison of HOMED with other single-cell guided methods across simulated and real datasets from PBMC, placenta, and lung tissues. For each tissue, mean MSE and mean PCC were obtained by averaging cell-type-specific deconvolution performance. Lower MSE and higher PCC indicate superior deconvolution accuracy. The best-performing metric for each tissue is highlighted in bold.

| Dataset | Method | Mean RMSE ↓ | Mean PCC ↑ |
| --- | --- | --- | --- |
| Simulated PBMC | HOMED | **0.01** | **1.00** |
|  | EpiSCORE | 0.25 | 0.64 |
|  | scDeconv | 0.13 | 0.74 |
|  | scMD | 0.15 | 0.77 |
| Simulated Lung | HOMED | **0.07** | **0.92** |
|  | EpiSCORE | 0.18 | 0.77 |
|  | scDeconv | 0.19 | 0.82 |
|  | scMD | 0.12 | 0.87 |
| Simulated Placenta | HOMED | **0.03** | **0.93** |
|  | EpiSCORE | 0.18 | 0.60 |
|  | scDeconv | 0.07 | 0.17 |
|  | scMD | 0.09 | 0.69 |
| Real PBMC | HOMED | **0.04** | **0.77** |
|  | EpiSCORE | 0.19 | 0.37 |
|  | scDeconv | 0.11 | 0.25 |
|  | scMD | 0.11 | 0.33 |
| Real Lung | HOMED | **0.12** | **0.69** |
|  | EpiSCORE | 0.20 | 0.49 |
|  | scDeconv | 0.20 | 0.67 |
|  | scMD | 0.17 | 0.69 |
| Real Placenta | HOMED | **0.06** | **0.41** |
|  | EpiSCORE | 0.13 | -0.10 |
|  | scDeconv | 0.14 | -0.26 |
|  | scMD | 0.16 | -0.02 |

