## Supplementary figures and images for "HOMED enables hierarchical and multimodal optimization of DNA methylation deconvolution across tissues" (from another version of this preprint: version 1, 2026-06-10)

### Supplemental Figure 1

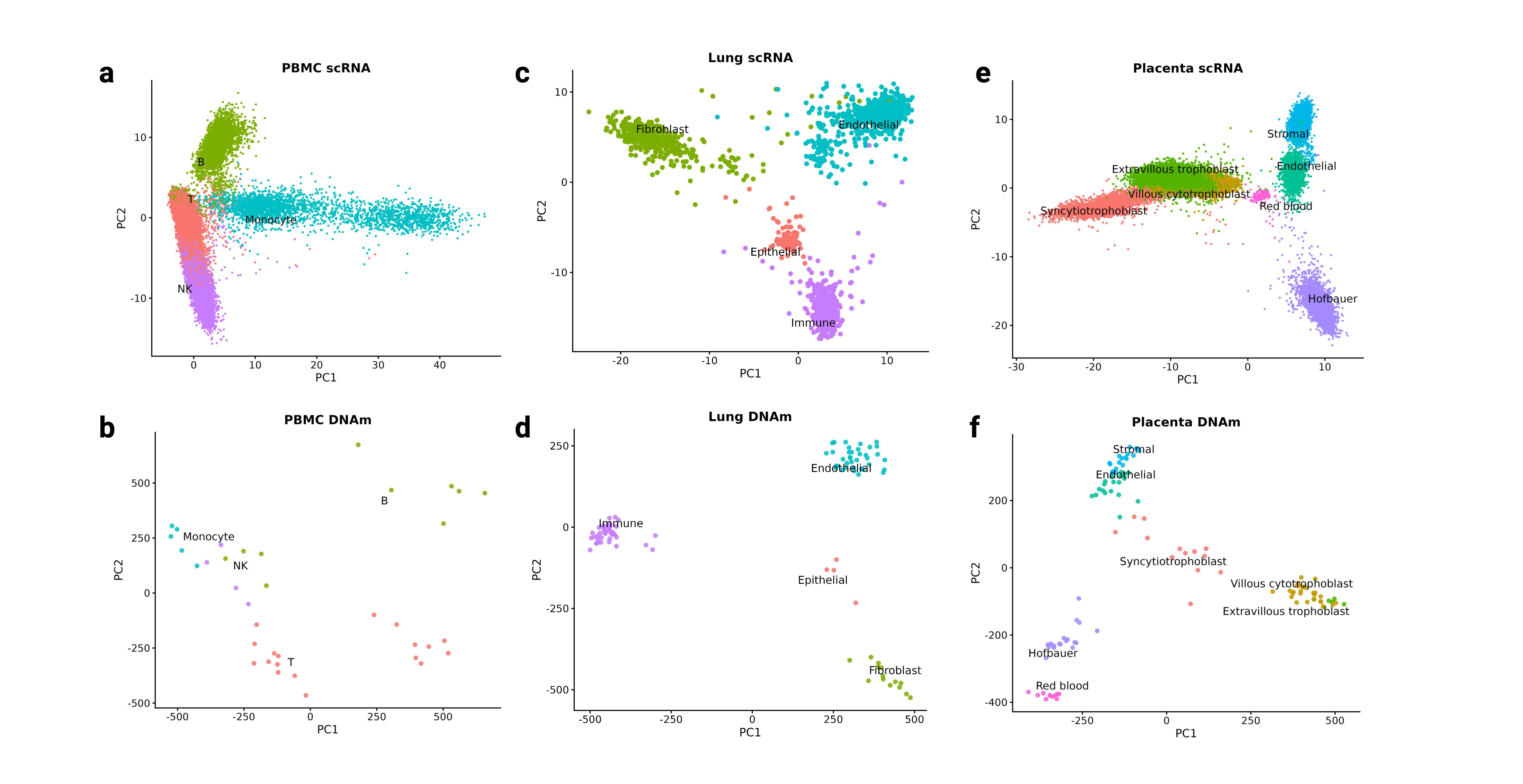

### Supplemental Figure 2

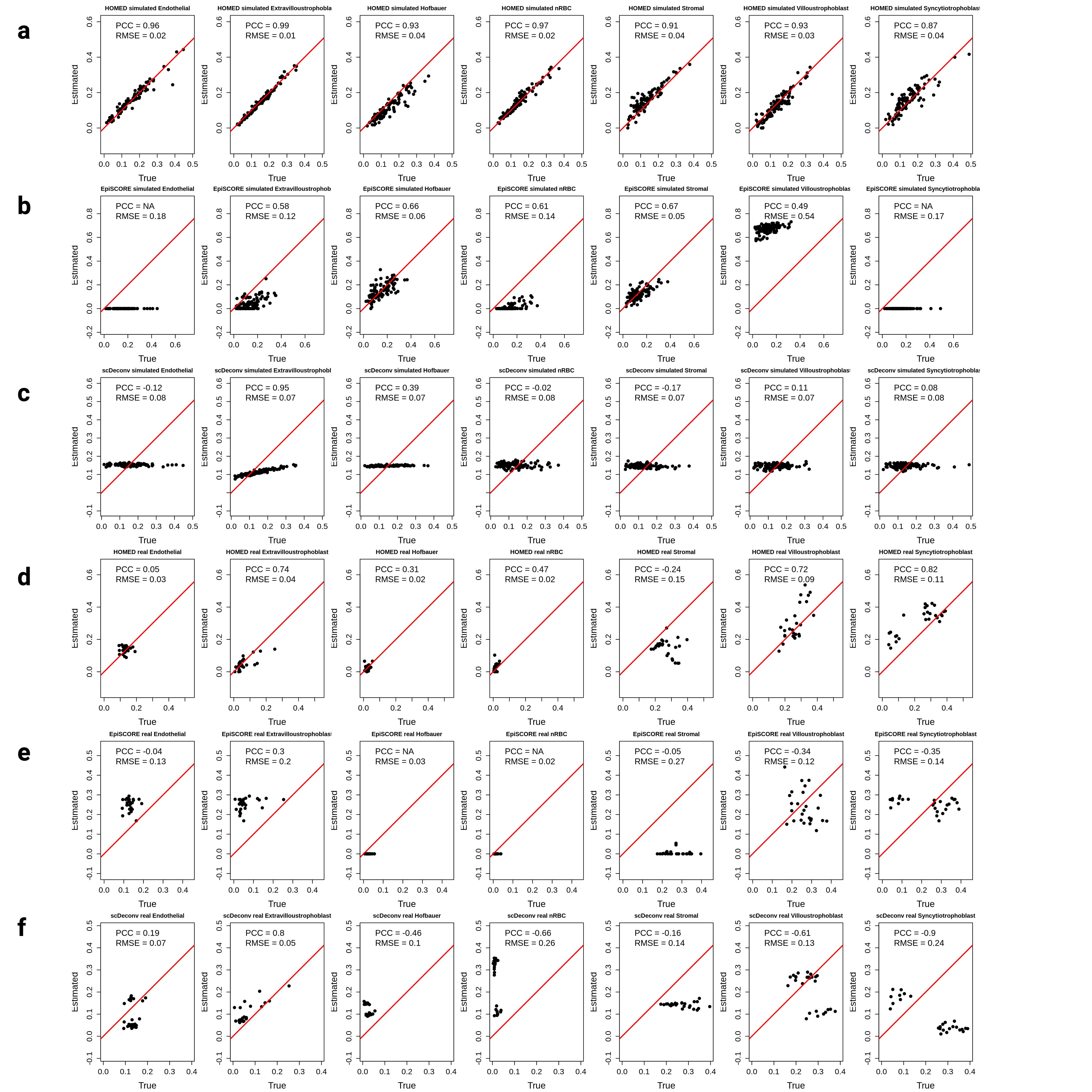

### Supplemental Figure 3

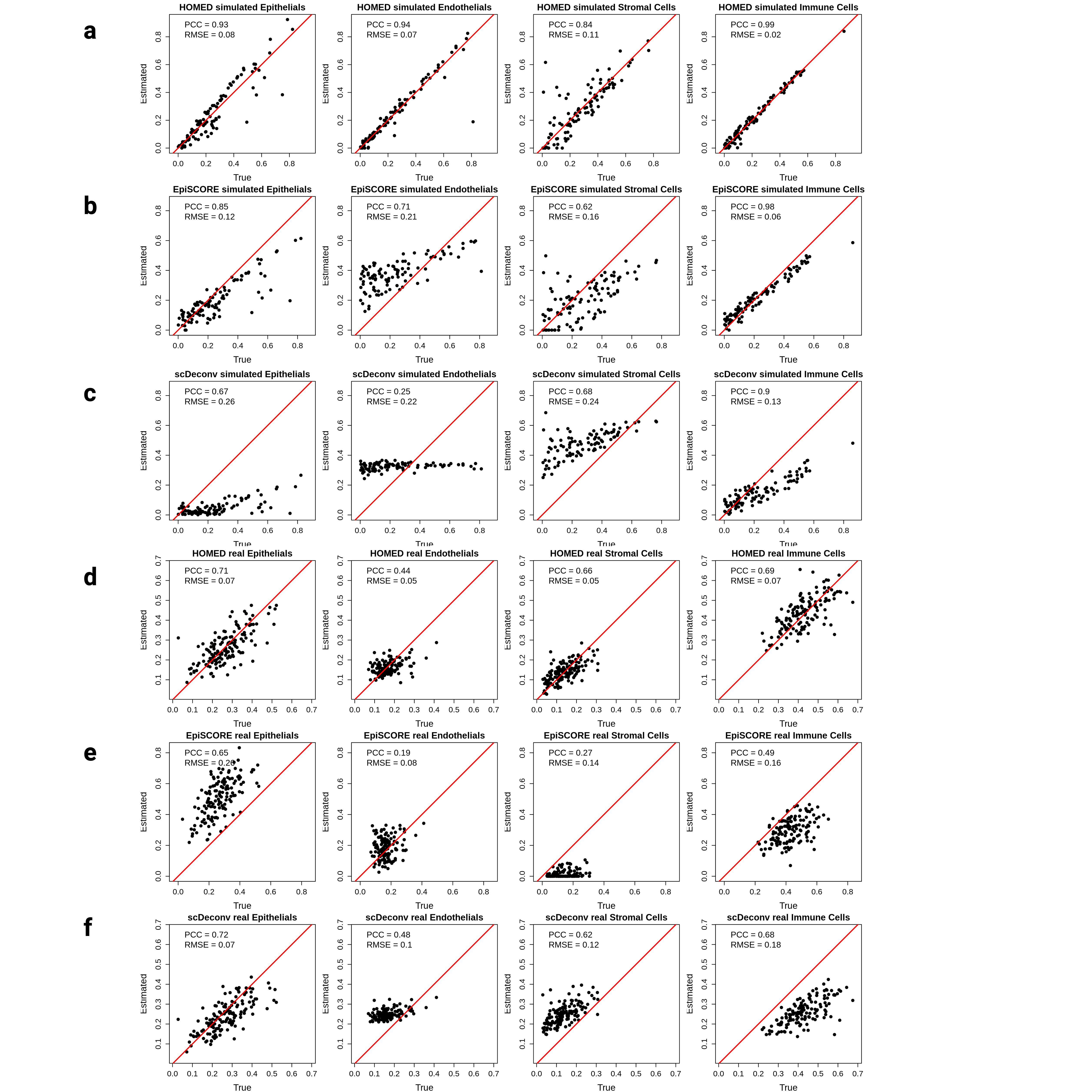
